# Noncanonical amino acid incorporation enables minimally disruptive labeling of stress granule and TDP-43 proteinopathy

**DOI:** 10.1101/2025.10.17.683020

**Authors:** Hao Chen, Haocheng Wang, Yuning Lu, Peng Chen, Zhongfan Zheng, Tao Zhang, Jiou Wang

## Abstract

We report a minimally disruptive labeling strategy for stress granule protein, G3BP Stress Granule Assembly Factor 1 (G3BP1), and ALS-linked protein, TAR DNA-binding protein 43 (TDP-43), using the fluorescent noncanonical amino acid Anap. By integrating genetic code expansion (GCE) with rational site selection, we achieved precise incorporation of Anap that preserves protein structure and function. In live cells and neurons, Anap labeling faithfully recapitulated localization, stress-induced dynamics, and recovery behavior, outperforming conventional fluorescent tags and enabling physiologically relevant visualization of protein pathobiology.

## Introduction

Fluorescent protein labeling remains a cornerstone of live-cell biology, yet conventional techniques rely heavily on large fusion tags, such as auto-fluorescent tags (AFPs) or small-molecule-binding motifs^1^, at limited positions (typically N-or C-terminal). These tags might potentially affect the structure, function, and even localization pattern of proteins, limiting their use for studying proteins with complex dynamics^2^. Alternatively, genetic code expansion (GCE) has emerged as a versatile labeling strategy to label proteins site-specifically in a minimally disruptive manner^3-8^.

GCE employs an engineered orthogonal aminoacyl tRNA synthetase/tRNA pair to incorporate non-canonical amino acids (ncAAs) at desired positions of proteins according to reassigned codons, most commonly the amber stop codon (TAG), thereby introducing a single-residue substitution within the protein of interest. Among these ncAAs, L-Anap (3-(6-acetylnaphthalen-2-ylamino)-2-aminopropanoic acid) is especially attractive for live-cell imaging. Anap is intrinsically fluorescent, exhibits polarity-sensitive emission spectra, and requires no post-incorporation modification^2, 5^. Despite these advantages, GCE-based Anap labeling has rarely been systematically applied to track disease-relevant protein dynamics in live mammalian cells. In this study, we developed an Anap-based labeling platform optimized for minimally disruptive labeling of two important proteins, G3BP1 and TDP-43, involved in membraneless organelles and neurodegenerative diseases, such as amyotrophic lateral sclerosis (ALS) and frontotemporal dementia (FTD).

G3BP1 is a core protein in stress granules, dynamic membraneless organelles related to stress response^9^. Altered stress-granule dynamics have been associated with ALS/FTD^10, 11^; however, whether stress granules directly drive neurodegeneration remains debated, as several studies suggest that stress granules primarily function as protective stress responses^12^. TDP-43 cytoplasmic inclusion is a hallmark of ALS/FTD pathology^13^ and is closely associated with dysregulation of RNA metabolism, ultimately leading to cellular defects^14^. Conventional fluorescent protein tags have enabled visualization of TDP-43 and G3BP1 in living cells; however, these approaches can perturb the native biophysical properties of the proteins being studied. For example, GFP or other fluorescently tagged TDP-43 usually requires additional modifications, such as deletion of the nuclear localization signal (NLS)^15, 16^, to induce cytoplasmic inclusion formation. Such manipulations introduce non-physiological conditions that may alter the native trafficking and aggregation behavior of TDP-43. As for G3BP1, tags like GFP may also cause unexpected effects on the phase separation or other dynamics of the protein. In contrast, Anap-based GCE strategy allows the minimally perturbative labeling and visualization of protein localization and stress-induced redistribution while preserving native protein architecture and function of both proteins. Importantly, the approach provides a generalizable genetically encoded platform for quantitatively examining the behavior of ALS-associated proteins in living cells. By enabling faithful monitoring of protein trafficking and stress-granule dynamics without extensive protein engineering, Anap-based GCE can offer a powerful strategy for probing molecular-scale mechanisms underlying ALS-linked proteinopathies.

## Results and Discussion

To implement the site-specific Anap incorporation system, we selected and generated two amber mutants, G3BP1^F337TAG^ and TDP-43^V100TAG^, using a combination of structural and functional criteria: exclusion from functional domains or localization signals, absence of disease-associated mutations, lack of post-translational modification, and low predicted structural impact by AlphaFold models. For G3BP1, the selected site was chosen to minimize interference with domains important for stress granule assembly, RNA binding, and protein-protein interactions. For TDP-43, the incorporation site was selected to avoid the major functional domains involved in RNA binding, nuclear localization, and aggregation-related behavior, thereby reducing the likelihood that Anap incorporation would perturb its native trafficking or function. More generally, we aimed to place the ncAA at positions likely to be solvent-accessible and tolerant of substitution, while avoiding highly conserved or functionally essential residues. Incorporation of Anap was achieved via co-expression of an orthogonal tRNA/synthetase pair in cells. To ensure that the fluorescence signal observed in our experiments was specifically derived from site-specific Anap incorporation rather than background fluorescence, we performed three control conditions. Specifically, we tested: (1) cells cultured with the addition of Anap, (2) cells expressing the Anap incorporation system with the addition of Anap, and (3) cells expressing both the TAG-mutated protein plasmid and the Anap incorporation system but without the addition of Anap. These control experiments were performed for both TDP-43 and G3BP1, and no observable fluorescence signal was detected under any of these conditions (Supplementary Fig. 1).

We first tested the feasibility of the Anap labeling system for G3BP1. In HeLa cells, G3BP1-Anap localized diffusely in the cytoplasm under basal conditions, closely matching antibody staining. Interestingly, a nuclear signal was detected with Anap but not with antibody, indicating the presence of nuclear pools of G3BP1 inaccessible to antibody detection. Upon sodium arsenite treatment, both the Anap and antibody signals colocalized within stress granules (Fig. 1A), validating the ability of Anap labeling to visualize the dynamics of G3BP1-driven stress granule formation. In addition, to independently validate protein expression, we performed western blot analysis in a G3BP knockout U2OS cell line, confirming the expression of G3BP1-Anap (Fig. 1H). To assess labeling fidelity, we compared the performance of G3BP1-Anap with G3BP1-GFP using fluorescence recovery after photobleaching (FRAP). Following stress, both labeled proteins localized to granules, but G3BP1-Anap exhibited significantly higher fluorescence recovery (∼53%) than G3BP1-GFP (∼33%) (Fig. 1C, 1F). These results suggest that G3BP1-Anap displays higher mobility compared with G3BP1-GFP, indicating that Anap labeling may provide a less perturbative approach for monitoring G3BP1 dynamics. Additionally, we examined the colocalization of G3BP1-Anap with TIA-1, another established stress granule marker. Under stress conditions, G3BP1-Anap largely colocalized with TIA-1 within stress granules. Interestingly, under basal conditions, the nuclear signal of G3BP1-Anap, which was not detected by antibody staining, appeared to partially colocalize with TIA-1 in several condensate-like structures. (Fig. 1I).

**Fig. 1:**
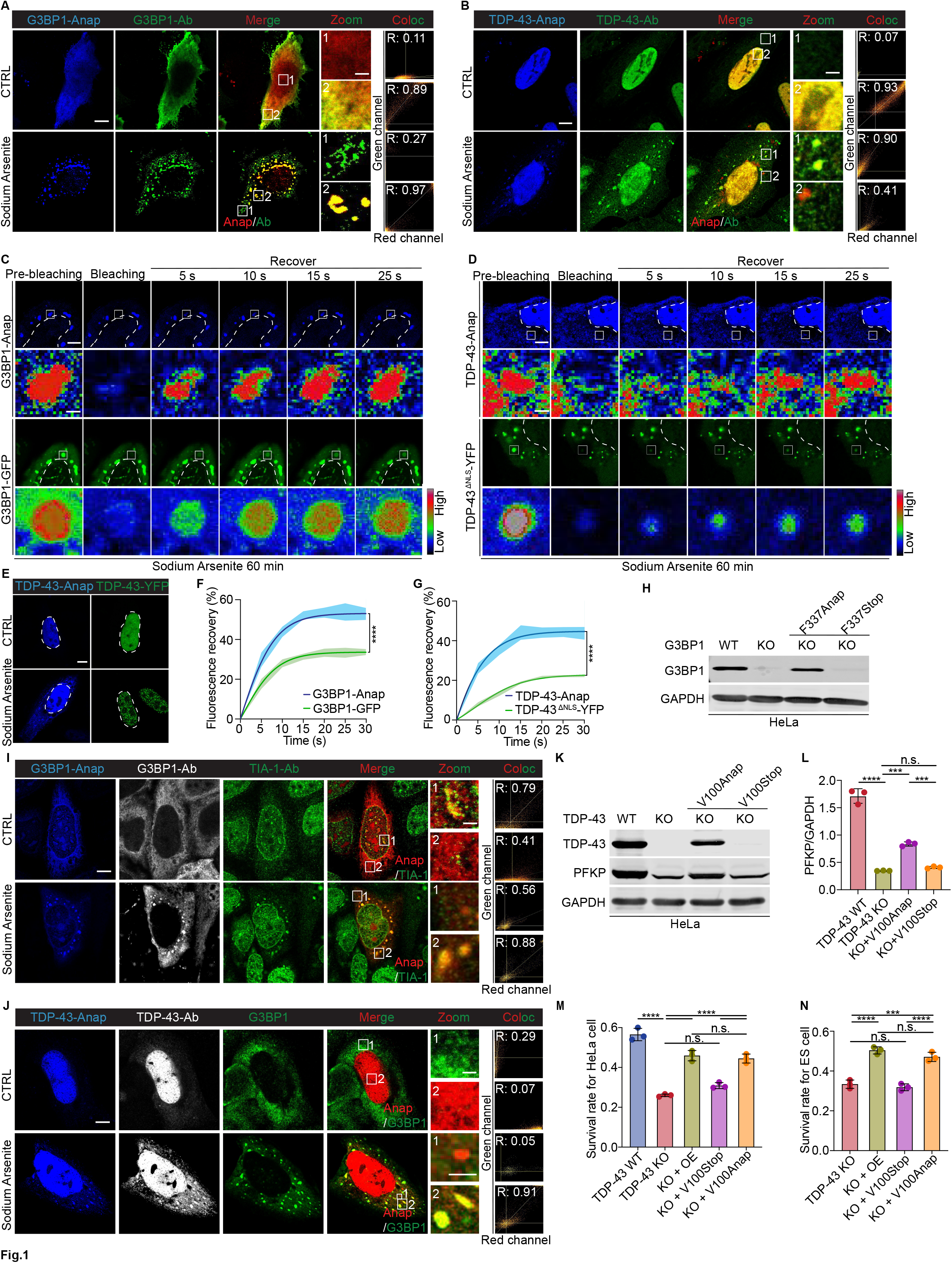
Anap-based labeling enables visualization of TDP-43 and G3BP1. A, B. HeLa cells expressing G3BP1-Anap and TDP-43-Anap under basal conditions or 250 μM sodium arsenite treatment. Anap and antibody signals are shown in blue and green, respectively; for merged channels, Anap was pseudo-colored red. Scale bars: 10 µm (overview), 3 µm (zoom). C, D. FRAP of G3BP1-Anap, G3BP1-GFP, and TDP-43-Anap, following 250 μM sodium arsenite treatment. One granule from each of three independent cells was selected and photobleached for FRAP analysis. ROI signal intensities are displayed in rainbow RGB (red-high, blue-low). Scale bars: 5 µm (cells), 1 µm (ROI). E. Comparison of HeLa cells expressing TDP-43-Anap and TDP-43-YFP under basal conditions or 250 μM sodium arsenite treatment. Here, Anap labeling and YFP labeling yield a blue signal and a yellow to green signal, respectively. F, G. Relative fluorescence recovery of each time point after photobleaching for G3BP1-Anap, G3BP1-GFP, TDP-43-Anap and TDP-43 ^ΔNLS^-YFP. Here, error bars with filled area are used for data presentation. FRAP recovery curves were compared using two-way ANOVA. H. Immunoblotting of wild-type and Anap labeled G3BP1. The mouse anti-G3BP1 antibody was used. KO: G3BP knock out; F337Stop/Anap: the expression of G3BP1^F337TAG^ via Anap labeling without or with the addition of Anap. I. Colocalization of G3BP1-Anap with TIA-1. The mouse anti-G3BP1 antibody and rabbit anti-TIA-1 antibody were used. Anap, G3BP1-Ab, and TIA-1-Ab signals are shown in blue, grey and green, respectively; for merged channels, Anap was pseudo-colored red. Scale bars: 10 µm (overview), 3 µm (zoom). J. Colocalization of TDP-43-Anap with G3BP1. The mouse anti-G3BP1 antibody and rabbit anti-TDP-43 antibody were used. Anap, TDP-43-Ab, and G3BP1-Ab signals are shown in blue, grey, and green, respectively; for merged channels, Anap was pseudo-colored red. Scale bars: 10 µm (overview), 3 µm (zoom). K. Immunoblotting of wild-type TDP-43, TDP-43-Anap and PFKP proteins in TDP-43 knockout HeLa cells. The Rabbit anti-TDP-43 antibody and rabbit anti-PFKP antibody were used. KO: TDP-43 knockout; V100Stop/Anap: the expression of TDP-43^V100TAG^ via Anap labeling without or with the addition of Anap. L. The expression levels of PFKP in TDP-43 knockout HeLa cells expressing wide-type TDP-43 or TDP-43-Anap. One-way ANOVA was used to compare the levels among groups. M. Survival of TDP-43 knockout HeLa cells expressing TDP-43-Anap after treatment with 12.5 μM sodium arsenite for 24h. Calcien EM staining was used to quantify cell survival, and the relative survival rate was calculated as the ratio of sodium arsenite-treated to untreated cells for each group. OE: overexpression of TDP-43. One-way ANOVA was used to compare levels among groups. N. Survival of iTDPKO inducible mouse ES cells expressing TDP-43-Anap. Here, cell counting-Lite 2.0 Luminescent cell viability assay kit was used to detect the survival rate of ES cells. TDP-43 knockout was induced by 4-HT (300ng/ml) for 5 days. The relative survival rate= 4-HT induction/DMSO for each group. One-way ANOVA was used to compare levels among groups. Colocalization threshold analysis was performed in Fiji/ImageJ to calculate the Pearson correlation coefficient (R) for each region of interest (A, B, I, J). The X and Y axes represent the fluorescence intensity values of the red and green channels, respectively. When signals are colocalized, pixels with high intensity in one channel correspond to high intensity in the other, forming a diagonal distribution. In contrast, non-colocalized signals cluster along the axes. A higher R value indicates a greater degree of colocalization. Scale bar, 3 μm. All quantitative data (F, G, L, M, N) are shown as mean ± SEM. ***P = 0.0001; ****P < 0.0001; n.s., not significant.

We next applied Anap labeling to TDP-43. Under basal conditions, the signal of TDP-43-Anap overlapped with that of anti-TDP-43 antibody staining, predominantly within the nucleus (Fig. 1B). Following sodium arsenite treatment, TDP-43-Anap mislocalized mostly to the cytoplasmic inclusions stained by TDP-43 antibody, with noticeable difference that a few puncta showed Anap signal only, and antibody gave more dispersed signal. By contrast, TDP-43-YFP failed to recapitulate this cytoplasmic mislocalization, instead forming prominent nuclear puncta under stress conditions (Fig. 1E), suggesting that large C-terminal tags may distort native localization of the protein. We then used YFP-tagged NLS-deleted TDP-43 (TDP-43^ΔNLS^-YFP) as a reference and performed FRAP analysis to compare the mobility of TDP-43-Anap and TDP-43^ΔNLS^-YFP. Fluorescence recovery of TDP-43-Anap reached ∼45% within 20 s after photobleaching, consistent with liquid-like dynamics. In contrast, TDP-43^ΔNLS^-YFP showed only ∼22% recovery, suggesting more solid-like dynamics (Fig. 1D, 1G). These results are consistent with previous reports describing relatively immobile aggregates formed by TDP-43^ΔNLS16^and illustrate the advantage of Anap-based labeling, which preserves native protein properties and enables real-time assessment of protein dynamics without introducing disruptive mutations. To further investigate the relationship between TDP-43-Anap-positive cytoplasmic inclusions and stress granules, we performed co-immunostaining with a G3BP1 antibody. Under stress conditions, most TDP-43-Anap-positive cytoplasmic inclusions colocalized with G3BP1-positive stress granules. However, a small subset of puncta contained TDP-43-Anap but lacked detectable G3BP1, suggesting that not all mislocalized TDP-43 is incorporated into stress granules under oxidative stress. These observations raise the possibility that additional mechanisms may contribute to the formation of TDP-43-positive cytoplasmic puncta (Fig. 1J).

To determine whether TDP-43-Anap retains biological function, we expressed it in a TDP-43 knockout HeLa cell line (Fig. 1K) and tested cell viability under oxidative stress. Following 12.5 μM sodium arsenite treatment for 24 hours, the expression of TDP-43-Anap significantly rescued cell survival, reaching levels comparable to wild-type TDP-43 overexpression (45% vs 46%; Fig. 1M). We further validated this finding in a mouse embryonic stem (ES) cell model with an inducible TDP-43 knockout (iTDPKO). For mouse ES cells, TDP-43 KO alone was sufficient to induce cell death. Here, either wild-type TDP-43 or TDP-43-Anap was expressed in iTDPKO mouse ES cells, where TDP-43 was deleted following induction with 4-hydroxytamoxifen (4-HT). Expression of TDP-43-Anap restored ES cell viability nearly to wild-type TDP-43 (47% vs 51%; Fig. 1N). We also evaluated TDP-43-dependent RNA splicing activity by examining the expression of PFKP, a well-established target that undergoes cryptic exon inclusion upon loss of TDP-43 function^17^. As shown in Figures 1K and 1L, expression of TDP-43-Anap in TDP-43 knockout HeLa cells restored PFKP expression, indicating that the Anap-labeled protein retains functional RNA splicing activity. These results demonstrate that TDP-43-Anap is capable of functionally compensating for endogenous TDP-43, supporting that the incorporation of Anap does not substantially disrupt the protein’s biological function.

Furthermore, to extend Anap labeling to neuronal systems, we applied this approach to label both proteins in primary mouse cortical neurons. The neurons were co-stained with human-specific anti-TDP-43 or human-specific anti-G3BP1 antibodies. Under basal conditions, the signal of G3BP1-Anap colocalized with antibody staining in the cytoplasm and relocalized to stress granule upon sodium arsenite treatment (Fig. 2A, 2B). Notably, nuclear Anap signal was again observed in neurons, suggesting additional pools of G3BP1 not captured by antibody staining.

**Fig. 2:**
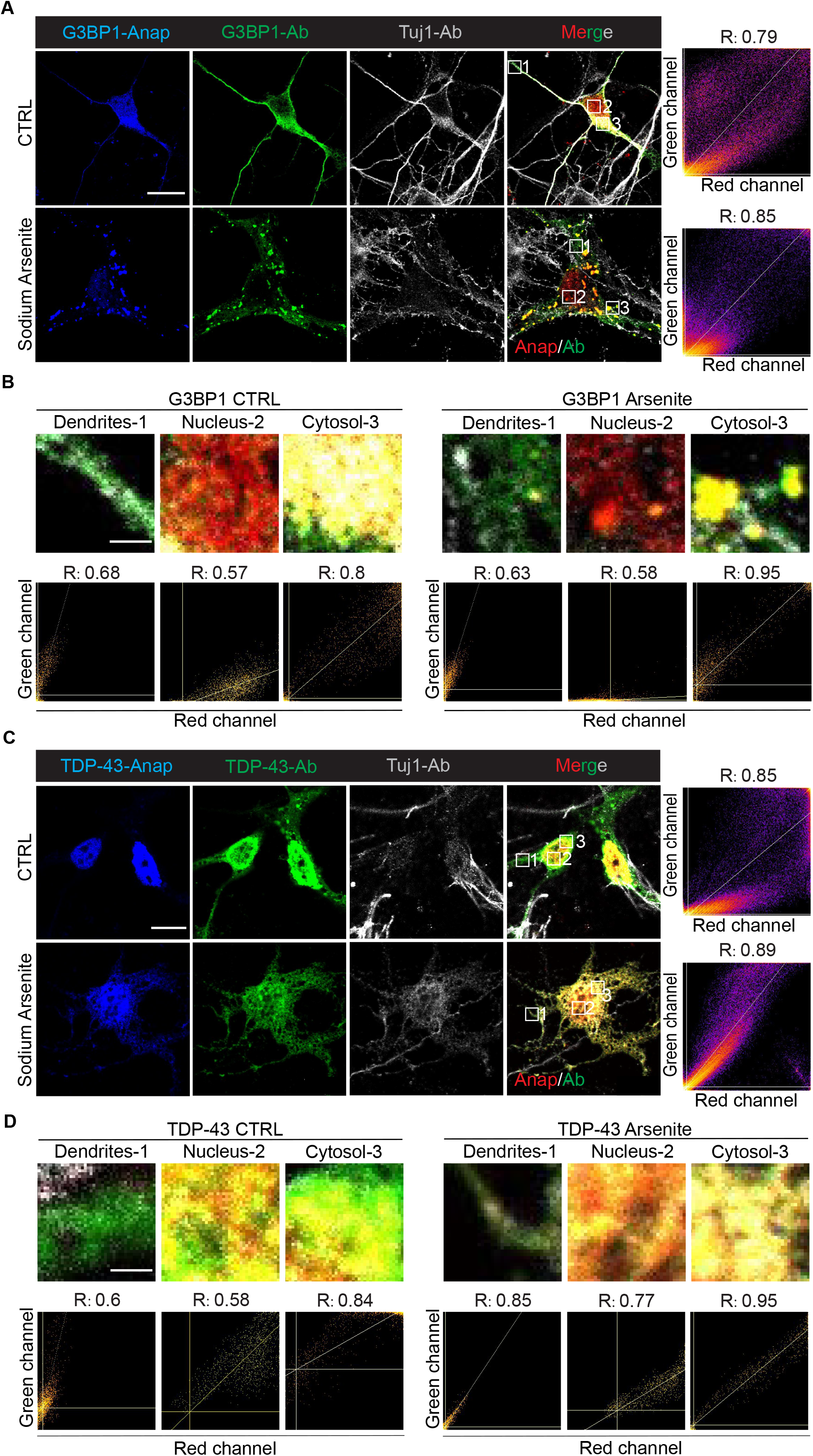
Anap labeling of TDP-43 and G3BP1 in neurons. A, C. Primary mouse cortical neurons expressing G3BP1-Anap and TDP-43-Anap under basal conditions or 250 μM sodium arsenite treatment. Cells were stained with anti-G3BP1 (human-specific) or anti-TDP-43 (human-specific) antibodies with chicken anti-Tuj1 as a neuron marker. Signals: Anap (blue, pseudo-colored red in merged images), antibody (green), Tuj1 (gray). Scale bar, 10 µm. B, D. The colocalization level of each region of interest for G3BP1 and TDP-43. Colocalization threshold analysis was performed in Fiji/ImageJ to calculate the Pearson correlation coefficient (R) for each region of interest. The X and Y axes represent the fluorescence intensity values of the red and green channels, respectively. When signals are colocalized, pixels with high intensity in one channel correspond to high intensity in the other, forming a diagonal distribution. In contrast, non-colocalized signals cluster along the axes. A higher R value indicates a greater degree of colocalization. Scale bar, 1 μm.

TDP-43-Anap localized to the neuronal nucleus under basal conditions and showed strong colocalization with anti-TDP-43 antibody. Interestingly, antibody staining appeared more diffusely cytoplasmic than Anap, suggesting improved signal specificity of Anap labeling with direct genetic incorporation. Under oxidative stress, both Anap and antibody signals colocalized within cytoplasmic inclusions (Fig. 2C, 2D). These findings demonstrated that the Anap system for G3BP1 and TDP-43 performs robustly in neuronal environments, a key setting for ALS/FTD research.

In summary, our results demonstrated that Anap-based GCE provides a minimally disruptive strategy for tracking the dynamic behavior of G3BP1 and TDP-43 in live cells (Fig. 3). In G3BP1, Anap faithfully reported stress granule assembly and preserved native mobility, unlike GFP fusions that impaired dynamics. In TDP-43, Anap labeling maintained nuclear localization under basal conditions and revealed liquid-like behavior of cytoplasmic inclusions during stress, in contrast to the aberrant nuclear puncta produced by YFP-tagged TDP-43. Critically, Anap-labeled TDP-43 retained biological activity, rescuing cell survival in TDP-43-deficient HeLa and stem cells. Moreover, we have generated stable TDP-43-Anap cell lines that exhibited consistent expression, protein localization, and stress-induced aggregation, providing stable cell models for TDP-43 research.

**Fig. 3:**
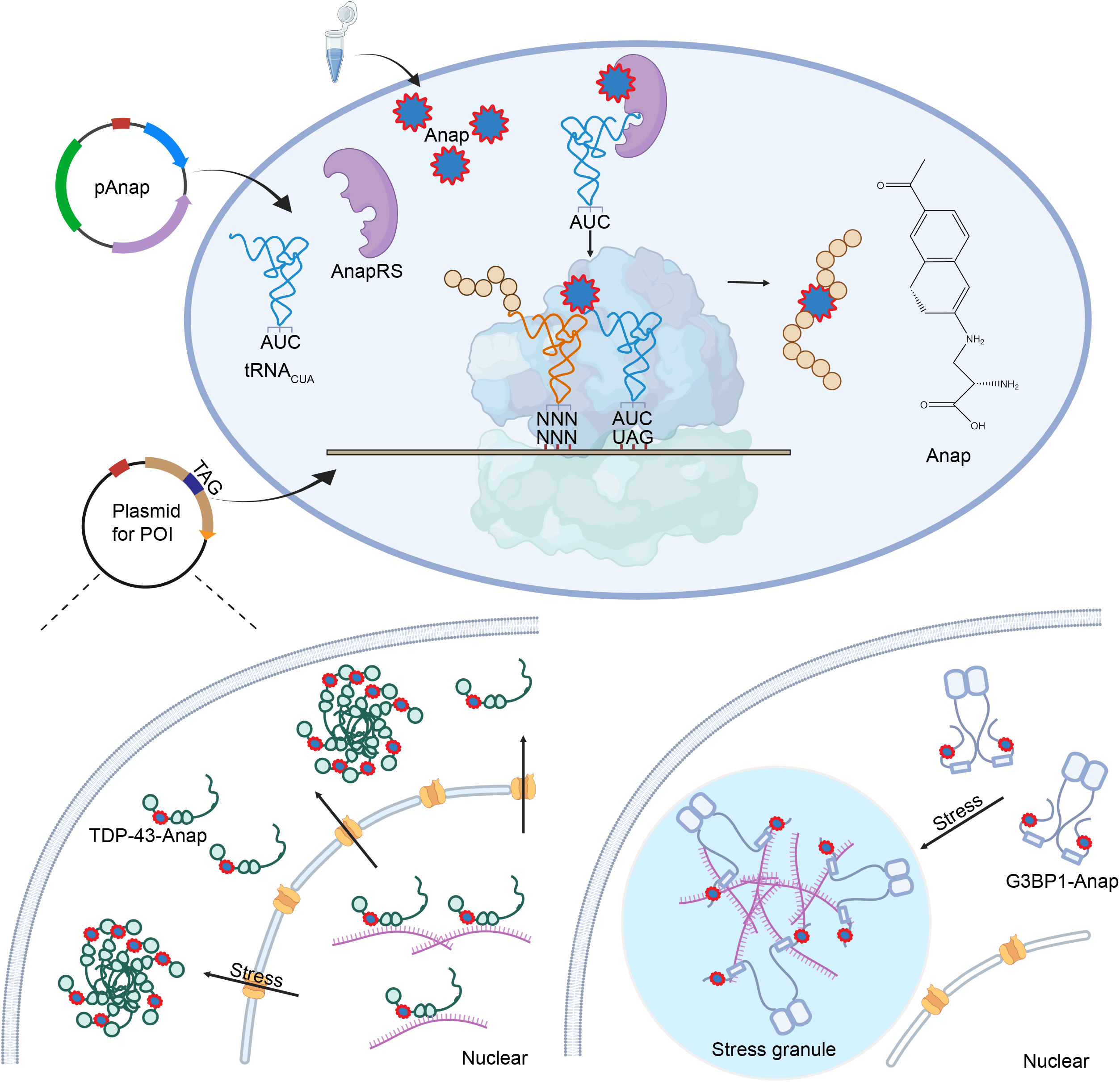
Schematic of the Anap labeling system for G3BP1 and TDP-43 using genetic code expansion. Briefly, two plasmids were required to express the protein with site-specific Anap incorporation, one for Anap incorporation and one for the mutated protein of interest (TAG introduction). As the plasmids were transfected into cells, the orthogonal Anap-tRNA synthetase would charge the Anap, a fluorescent amino acid, to its cognate tRNA, and the tRNA would incorporate the Anap site-specifically into the protein of interest in response to the TAG stop codon. Cells expressing Anap-labeled TDP-43 and G3BP1 were subsequently imaged by confocal microscopy, either after fixation or in live-cell conditions. Upon stress conditions, TDP-43-Anap redistributes to the cytoplasmic inclusions, and G3BP1-Anap assembles into stress granule.

By enabling high-fidelity visualization of both stress granule dynamics and TDP-43 aggregation in live cells and primary neurons, Anap labeling bridges a critical gap between structural preservation and functional readout. The ability to monitor native protein behavior without perturbation provides a unique opportunity to study early events in ALS/FTD progression, such as stress granule maturation, protein cytoplasmic mislocalization, and aggregate fluidity, at a resolution inaccessible with conventional tagging approaches.

## Acknowledgements

This work was supported by the National Institute of Health (NIH) (NS074324, NS089616, NS110098, and NS128494) and the Walder Foundation. We thank Peter G. Schultz from Scripps Research for providing the pAnap plasmid, Shawn M. Ferguson from Yale University for the TDP-43 knockout HeLa cell line, J. Paul Taylor from St. Jude Children’s Research Hospital for the G3BP knockout U2OS cell line, and Philip Wong from Johns Hopkins University for the iTDPKO mouse ES cell line. We also thank the members of Wang’s lab for discussion and suggestions.

## Author contributions

H.C. performed and analyzed most of the experiments. H.C.W. helped with plasmid preparation, western blot, and imaging. Y.N.L. provides cells and helped with data preparation. P.C. provides mouse primary cortical neurons. Z.F.Z helped with imaging. T.Z. helped culture iTDPKO mouse ES cells. H.C. and J.W. designed the studies and wrote the paper with inputs from other authors.

## Methods and materials

### General information

Oligonucleotide synthesis was performed by IDT, and Sanger sequencing of DNA plasmids and PCR products was performed by Quintara. L-ANAP (trifluoroacetate salt) (15436) used in this study was purchased from Cayman. For immunostaining, the following primary and secondary antibodies were used: mouse anti-human G3BP (BD Biosciences, 611126), rabbit anti-TDP-43 (Proteintech, 10782-2-AP), mouse anti-TDP-43 (human-specific, monoclonal; Proteintech, 60019-2-Ig), Rabbit anti-human TIA-1 (MBL Life science, RN014P), chicken anti-βIII-tubulin (Tuj1; GeneTex, GTX85469), donkey anti-mouse Alexa Fluor 555 (Thermo Fisher, A-31570), donkey anti-rabbit Alexa Fluor 488 (Thermo Fisher, A-21202), donkey anti-chicken Alexa Fluor 647 (Thermo Fisher, A78952), and donkey anti-rabbit Alexa Fluor Plus 800 (Thermo Fisher, A32808).

### Plasmids

pAnap plasmid was a gift from Peter G. Schultz’s lab^2^. pCMV-G3BP1 was purchased from Sino Biologics. pRK5-TDP-43, pEGFP-G3BP1, and pCMV/TO-TDP-43-YFP were conducted by our lab. pLVX-Puro-TDP-43-WT was a gift from Shawn Ferguson’s lab^18^ and reconstructed into pLVX-Puro-TDP-43-ΔNLS. Site-directed mutagenesis was conducted using the NEB Q5 site-directed mutagenesis kit (NEB, E0554). TAG substitutions were introduced by PCR amplification with Q5 Hot Start High-Fidelity DNA Polymerase using primers designed with NEBaseChanger. Then the PCR products were incubated with an enzyme mix consisting of a kinase, a ligase, and DpnI to rapidly circularize the PCR products and remove the template DNA. And then the mix will be transformed into DH5α cells.

### Cell culture

Mammalian cell lines were cultured at 37°C and 5% CO_2_ in humidified incubators. HeLa cells (ATCC) were cultured in DMEM/F12 (Corning, 10-013-CV) supplemented with 10% FBS (Gibco, A5256801). The TDP-43 knockout HeLa cell line was a gift from Shawn M. Ferguson^18^ and the G3BP knockout U2OS cell line was a gift from J. Paul Taylor^9^. Primary cortical neurons were prepared as previously described ^19^. Briefly, the cortex of the mouse embryo was separated into HBSS on ice, then digested into single-cell suspension with 0.25% trypsin and 0.1mg/ml DNase I at 37°C for 20 min. Dissociated cells were washed twice and resuspended in plating media (DMEM supplemented with 10% FBS) before seeding onto poly-D-lysine (Gibco, A3890401)-coated plates. Cells were cultured for 3-4 hours in plating medium to settle down, and then the medium was replaced with maintenance medium (neurobasal medium (Gibco, 21103049)) supplemented with 2% B-27 supplement (Gibco,17504044), 1% GlutaMax (Gibco, 35050061), and 1% penicillin/streptomycin (Gibco, 15140163). The iTDPKO inducible mouse embryonic stem ES cell line was a gift from Philip Wong^20^. Cells were maintained on attachment factor (Gibco, S-006-100) coated plates in 2i media containing half of DMEM/F12 and half of Neurobasal media complemented with L-Glutamine (Gibco, 25030081), B-27 supplement, N2 supplement (Gibco, 17502048), BSA (Gibco, 15260037), 1% penicillin/streptomycin (Gibco, 15140163), PD0325901 (MedChemExpress, 391210-10-9), CHIR99021(MedChemExpress, 252917-06-9), monothioglycerol(Millipore Sigmal, M6145), and mLIF(Millipore Sigma, ESG1107).

### Transfection

HeLa cells were transfected using Lipofectamine 2000 reagent (ThermoFisher, 11668030). Cells were seeded 24 h before transfection, and plasmid DNA was mixed with Lipofectamine 2000 at a ratio of 1 µg DNA:2 µL reagent in Opti-MEM medium (Thermo Fisher, 31985070). After 15 min incubation at room temperature, the complexes were added to the cells. Six hours post-transfection, the medium was replaced with fresh growth medium supplemented with or without 20 µM L-Anap, and cells were incubated overnight before further experiments.

Mouse primary cortical neurons and inducible TDP-43 knockout (iTDPKO) mouse ES cells were transfected using Lipofectamine 3000 (Thermo Fisher, L3000015) according to the manufacturer’s instructions. For neurons, first, the total DNA was diluted and mixed with p3000 reagent in Opti-MEM media, and the Lipofectamine 3000 was diluted in Opti-MEM media separately. Second, both diluted DNA and Lipofectamine 3000 were mixed and incubated at room temperature for 15 minutes. The final mix would then be added to cells seeded on poly-D-lysine-coated plates at DIV5. After incubation overnight, the media were half-replaced with fresh neurobasal media with 10µM Anap, and the neurons were incubated for two additional days (the media were half-replaced each day to keep the Anap supply). Differently, for mouse ES cells, the DNA-lipo3000 mix was added directly into the cell suspension, and the suspension was added to attachment factor-coated plate. After overnight incubation, the media for ES cells was replaced with fresh 2i media supplemented with 10 µM Anap and any further treatments.

### Anap labeling and immunofluorescence in cell fixation

6h post-transfection, the cells were cultured in medium supplemented with L-Anap for another 20-24 h. Cells were then washed three times with DPBS to remove excess Anap and incubated in fresh medium for 1 h before treatment with 250 µM sodium arsenite for anther 1h. After the treatment, the cells were washed with DPBS three times and then fixed with 4% paraformaldehyde (PFA; Millipore Sigma, 158127) for 15-20 mins at room temperature. Fixed cells were permeabilized and blocked in immunofluorescence blocking buffer (Cell Signaling Technology, 12411) containing 0.1% Triton X-100 (Millipore Sigma, X100) for 45 mins. Immunostaining was performed using anti-TDP-43 and anti-G3BP1 antibodies in both HeLa cells and primary cortical neurons, with anti-Tuj1 antibody included as a neuronal marker. Images were acquired using a Leica SP8 confocal microscope.

### Anap labeling and GFP/YFP tagging in live-cell imaging

For Anap labeling, HeLa cells were seeded on 35-mm glass-bottom dishes (Cellvis, D35-20-1.5H), transfected, and incubated with L-Anap following the same protocol used for fixed-cell imaging. After removal of excess Anap, cells were cultured in fresh medium for 2-3 h before live cell imaging. For GFP/YFP tagging, cells expressing the tagged proteins were directly ready for live cell imaging. Before FRAP, cells expressing Anap-labeled or GFP/YFP-tagged TDP-43 or G3BP1 were treated with 250 μM sodium arsenite for 1 h. Next, regions of interest (ROIs) corresponding to protein signals were selected for FRAP analysis using a Leica SP8 confocal microscope.

### Cell survival tests

Cell viability was assessed using Calcein AM staining (Invitrogen, C1430). HeLa cells were treated with 12.5 µM sodium arsenite for 24 h, followed by incubation with 3 µM Calcein AM.

Fluorescence from viable cells was measured using a Synergy H1 Hybrid Multi-Mode Plate Reader (BioTek) at excitation/emission wavelengths of 485/535 nm.

For mouse ES cells, the conditional knockout of TDP-43 was induced by 4-hydroxytamoxifen (4-HT, Millipore Sigma, H7904-5MG) treatment (300ng/ml) for 5 days. Cell viability was assessed using the Cell Counting-Lite 2.0 Luminescent Cell Viability Assay Kit (Vazyme, DD1101-02).

### Immunoblotting

Cells were washed three times with PBS and lysed in ice-cold RIPA buffer [50 mM Tris-HCl (PH 7.5), 150 mM NaCl, 1% NP40, 0.1% SDS, 100 mM NaF, 0.5% sodium deoxycholate, 17.5 mM beta-glycerophosphate, 1 mM PMSF, and protease inhibitor cocktail (1:200, Millipore Sigma, P8340)]. Lysates were sonicated and clarified by centrifugation (12,000 rpm, 20 min, 4 °C), and supernatants were collected. Protein concentrations were determined using the BCA assay (Thermo Fisher, Cat. No. 23225). Equal amounts of protein were resolved by SDS-PAGE and transferred to membranes, which were blocked with 5% BSA and incubated with primary antibodies overnight at 4 °C. After three washes with TBST, membranes were incubated with fluorescence-conjugated secondary antibody dilution at room temperature for 2h. Blots were scanned and imaged by LI-COR Odyssey M scanner.

### Image analysis and statistical analysis

All images were processed and analyzed by Fiji/ImageJ software, and the colocalization of signals was analyzed by the colocalization threshold analysis plugin in ImageJ. Experiments requiring quantification were repeated at least three times independently. The FRAP results were analyzed by ImageJ to quantify the fluorescence intensity at each time point. The average relative intensities (normalized to pre-bleaching intensity) of each time point were analyzed in Prism 10 software. For survival tests, the average relative rates (normalized to the control group) for each group were analyzed in Prism 10 software, and the one-way analysis of variance (ANOVA) was used to compare the significance between each group. All data were presented as means ± SEM.

## Figure legend

**Supplementary Fig. 1: Controls for Anap-based labeling of TDP-43 and G3BP1 in HeLa cells**

A. Controls for TDP-43-Anap labeling. Four experimental conditions were tested: (1) HeLa cells expressing both the TAG-mutated TDP-43 plasmid and the Anap incorporation system in the presence of Anap (TDP-43-Anap); (2) cells expressing both plasmids without Anap (TDP-43-Stop); (3) cells cultured with Anap only; and (4) cells expressing the Anap incorporation system with the addition of Anap. Signals for TDP-43-Anap, TDP-43 antibody staining, and G3BP1 antibody staining are shown in blue, grey, and green, respectively. The Anap signal is pseudo-colored red in the merged images. B. Controls for G3BP1-Anap labeling. The same four conditions were tested: (1) cells expressing the TAG-mutated G3BP1 plasmid and the Anap incorporation system in the presence of Anap (G3BP1-Anap); (2) cells expressing both plasmids without Anap (G3BP1-Stop); (3) cells cultured with Anap only (+ Anap only); and (4) cells expressing the Anap incorporation system with the addition of Anap (Anap system+Anap). Signals for G3BP1-Anap, G3BP1 antibody staining, and TIA-1 antibody staining are shown in blue, grey, and green, respectively. The Anap signal is pseudo-colored red in the merged images. Scale bars, 40 μm.

