## Supplementary figures and images for "Noncanonical amino acid incorporation enables minimally disruptive labeling of stress granule and TDP-43 proteinopathy"

### Supplementary Figure. 1

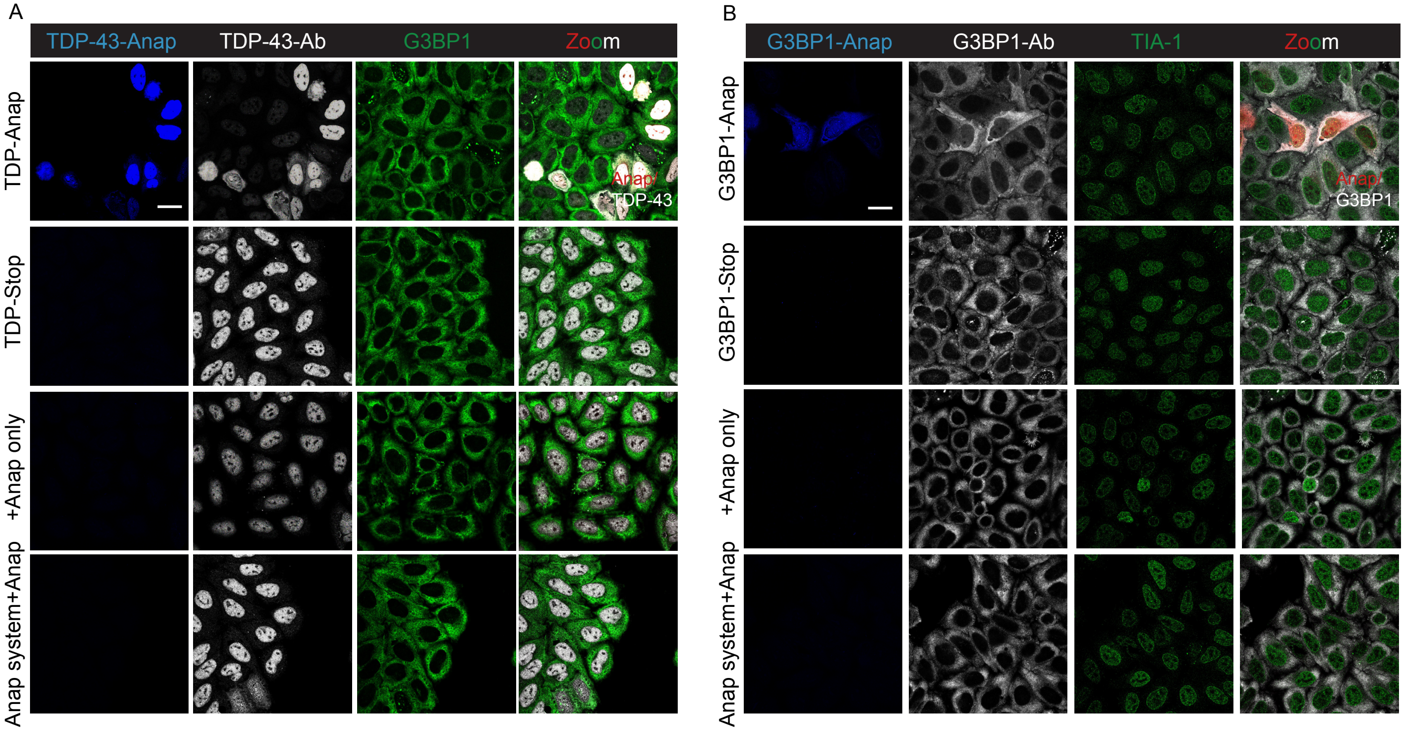


Supplementary fig. 1
